# Admixture mapping reveals loci for carcass mass in red deer x sika hybrids in Kintyre, Scotland

**DOI:** 10.1101/2021.04.29.442014

**Authors:** S. Eryn McFarlane, Josephine M. Pemberton

**Affiliations:** Institute of Evolutionary Biology, School of Biological Sciences, University of Edinburgh, Edinburgh, UK; Department of Biology, Lund University, Lund, Sweden

## Abstract

We deployed admixture mapping on a sample of 386 deer from a hybrid swarm between native red deer (*Cervus elaphus*) and introduced Japanese sika (*Cervus nippon*) sampled in Kintyre, Scotland to search for Quantitative Trait Loci (QTL) underpinning phenotypic differences between the species. These two species are highly diverged genetically (F_st_ between pure species, based on 50K SNPs, = 0.532) and phenotypically: pure red have on average twice the carcass mass of pure sika in our sample (38.7kg vs 19.1 kg). After controlling for sex, age and population genetic structure we found ten autosomal genomic locations with QTL for carcass mass. Effect sizes ranged from 0.191 to 1.839 Kg and as expected, in all cases the allele derived from sika conferred lower carcass mass. The sika population was fixed for all small carcass mass alleles, whereas the red deer population was typically polymorphic. GO term analysis of genes lying in the QTL regions are associated with oxygen transport. Although body mass is a likely target of selection, none of the SNPs marking QTL are introgressing faster or slower than expected in either direction.

## Introduction

To understand the relationship between genetic variation and phenotypic variation, and eventually the link between genetic variants and fitness, is a goal of evolutionary genetics. By understanding the genetic architecture of phenotypic traits, we can then ask how selection could act on a trait, make predictions of how a trait might change over time, or how the trait could respond to environmental change (Barton and Keightley 2002). In the context of hybridization, it is informative to understand the genetic architecture of the phenotypic traits that differ between hybridizing species. This is particularly relevant when human influences lead to increased hybridization (Grabenstein and Taylor 2018) and there is the potential for extinction via hybridization to decrease biodiversity (Rhymer and Simberloff 1996; Brennan et al. 2015; Todesco et al. 2016).

Genetic mapping in hybrid zones is particularly powerful because of the opportunity to use admixture mapping on recombinant individuals (Rieseberg and Buerkle 2002). The assumption of admixture mapping is that hybrid individuals have mosaic genomes that have been formed as the result of introgression, selection, and genetic drift (Buerkle and Lexer 2008; Winkler et al. 2010; Seldin et al. 2011). Coupled with divergent phenotypes, this allows for quantitative trait locus (QTL) mapping using fewer markers than are needed for typical genome wide association studies (Rieseberg and Buerkle 2002). Natural hybrid zones can be extremely powerful for detecting QTLs because the phenotypes of hybrids are often intermediate, (Buerkle and Lexer 2008). Admixture mapping is most powerful when both the phenotype and genotypes are divergent between the two parental populations and when there individuals are sampled across the ancestry and phenotype spectrum (Buerkle and Lexer 2008).

Admixture mapping has been used in human populations, wild plants and in some wild animals, but less so in wild mammals. Specifically, admixture mapping has been used extensively to find genes for disorders in human populations (Patterson et al. 2004; Smith et al. 2004; Shriner 2013), to search for genes related to reproductive isolation in *Populus* hybrid zones (Lexer et al. 2007; Lexer et al. 2010), and for morphological and phytochemical traits in these hybrid zones (Bresadola et al. 2019) and for traits such as plumage colour, migration behaviour and beak size in birds (Chaves et al. 2016; Delmore et al. 2016; Brelsford et al. 2017), melanoma and tail fin morphology in swordtail fish (*Xiphophorus malinche* and *X. birchmanni* (Powell et al. 2020; Powell et al. 2021), and wing pattern variation in butterflies (Lucas et al. 2018). In wild mammal systems, admixture mapping has been used to discover 10 genomic regions for craniofacial shape variation and 23 single nucleotide polymorphisms (SNPs) associated with leg bone length in mice (*Mus musculus musculus x M. m. domesticus;* (Pallares et al. 2014; Škrabar et al. 2018), and to associate introgressed genomic regions with body size and skeletal growth in coyotes and wolves (*Canus latrans and C. lupus*; (vonHoldt et al. 2016)). While admixture mapping is suitable for gene mapping in wild mammals, the best systems would be hybrid swarms with substantial variation in focal phenotypes.

Anthropogenic hybridization between red deer (*Cervus elaphus*) and sika (*C. nippon*) in Scotland (Senn et al. 2009, McFarlane et al. 2020), offers an opportunity to use admixture mapping to identify the genetic architecture of an extremely variable phenotype, in this case, carcass mass. Briefly, sika were introduced to Scotland in the 19^th^ century, and hybrid individuals in Kintyre are common (McFarlane et al. 2020). Carcass mass of red deer males in Argyll ranges between 55 and 106 kg, while carcass mass of red deer females ranges from 51-61 kg; by comparison sika in Scotland have an average carcass mass of 30 kg (males) and 24 kg (females; (Harris and Yalden 2008) indicating substantial divergence in this trait between the two species. Hybrid individuals have intermediate phenotypes correlated with their admixture proportion (Senn et al. 2010). Carcass mass is the weight in kilograms of the animal at death, following the removal of the head, internal organs, lower legs and blood. Thus, carcass mass is approximately 60-70% of live mass (Mitchell and Crisp 1981). Red deer and sika in Scotland are quite genetically diverged, with a genome wide F_st_ of 0.532 (95% confidence interval: 0.529 – 0.534 McFarlane et al. 2020a), although it should be noted that there is substantial variation in divergence across the genome(McFarlane et al. 2021). If there is a SNP for carcass mass that is in a causal region, or in linkage disequilibrium with a causal region, we should have high power to detect it, based on the F_st_ between red deer, the large phenotypic divergence and the estimated number of generations since admixture began (approximately 6-7; (Crawford and Nielsen 2013; McFarlane et al. 2020).

The goals of this study are to use the red-sika hybrid system to 1) identify any large effect QTL for carcass mass, 2) Inspect the direction of effect of any QTL found, with the prediction that the sika-specific allele will confer lower mass and 3) search for nearby genes and analyze their putative functions. In general we expect carcass mass in deer to have a polygenic architecture, as is the case in morphological traits in several wild systems (e.g. Soay sheep (Bérénos et al. 2015), collared flycatchers and house sparrows (Silva et al. 2017), and great tits (Santure et al. 2013; Santure et al. 2015). However, there could also be some large effect QTL, as have been found for human height (Yang et al. 2010), cattle (Bhuiyan et al. 2018; Roberts 2018; Pegolo et al. 2020), and such QTL are particularly likely in an admixed population.

## Methods

We analyzed 513 deer samples collected from 15 forestry sites in the Kintyre region of Scotland between 2006 and 2011. The Forestry Commission Scotland (now Forestry and Land Scotland) culled the deer as part of normal deer control measures, in which animals were shot as encountered, regardless of phenotype or suspected species (Smith et al. 2018). Ear tissue samples were stored in 95% ethanol, and animals were sexed, aged (from tooth eruption and wear) and weighed to the nearest kg within 24 hours of harvest (Senn and Pemberton 2009). Of the 513 deer sampled and genotyped, carcass mass was available for 386 animals.

### DNA extraction and SNP Genotyping

The deer were genotyped on the Cervine Illumina iSelect HD Custom BeadChip, which has 53,000 attempted SNP assays, using an iScan instrument (Huisman et al. 2016). When this SNP chip was developed, SNPs were selected to be spaced evenly throughout the genome based on the bovine genome with which the deer genome has high homology, although we use the deer linkage map in the present study (Johnston et al. 2017). The majority of SNPs were selected because they were polymorphic in red deer, specifically those red deer that are part of a long term monitoring project on the Isle of Rum, but 4500 SNPs were also selected to be diagnostic between either red deer and sika or red deer and wapiti (*Cervus canadensis*) (Brauning et al. 2015).

We used the DNeasy Blood and Tissue Kit (Qiagen) according to the manufacturer’s instructions to extract DNA for SNP analysis, with the exception that we eluted twice in 50μl buffer TE to obtain DNA at a sufficiently high concentration. We assayed the concentration of extractions using the Qubit™ dsDNA BR Assay Kit (Invitrogen). If an extraction was below 50 ng/μl, it was vacuum-concentrated, re-extracted or omitted from SNP analysis. Each 96 well plate had a positive control, and genotypes were scored using the clusters from a previous study (Huisman et al. 2016; McFarlane et al. 2020).

We followed the same protocol as McFarlane et al. (2020) for quality control, and to estimate the proportion of red deer ancestry for each individual (Q score). We used PLINK for all quality control (Purcell et al. 2007). Specifically, we excluded individual samples with a call rate of less than 0.90, deleted loci with a minor allele frequency of less than 0.001 and/or a call rate of less than 0.90 (McFarlane et al. 2020), but we did not exclude SNPs based on Hardy Weinberg Equilibrium (HWE) as admixed samples are not expected to be in HWE. To assign a Q score to each individual we used ADMIXTURE (Alexander et al. 2009). If the credible interval around the Q score overlapped 0, an individual was considered pure sika, if the CI overlapped 1 then it was pure red deer and the individual was considered a hybrid if the CIs overlapped neither 0 or 1 (McFarlane et al. 2020).

### Admixture mapping

We used Bayesian sparse linear mixed models (BSLMMs) in *gemma* for admixture mapping (Zhou et al. 2013). BSLMMs model the genetic architecture of traits while controlling for relatedness, thus giving an estimate of the proportion of phenotypic variance explained by combined effects of polygenic and large effect SNPs. SNP effects are drawn from two distributions, one distribution where it is assumed that all SNPs have a small to negligible effect, and a second distribution where some SNPs are assumed to have a larger effect drawn from a different distribution (i.e. the sparse effects; Zhou et al. 2013). BSLMMs include a kinship matrix to account for phenotypic similarity based on overall relatedness or genetic similarity. Inclusion of this kinship matrix removes the effect of population structure when determining whether individual SNPs have a significant effect on the trait (Zhou et al. 2013). From the BSLMM models, we can extract estimates of the proportion of variance in the phenotype explained (PVE) by the sparse effects and the random effects, as well as the proportion of the genetic variance explained (PGE) by the sparse effects. The product of PVE and PGE is the proportion of phenotypic variance explained by the sparse effects, known as the narrow sense heritability (h^2^).

A BSLMM cannot be run with a covariate matrix, although covariates can be included as additional SNP effects using the command ‘--not-snp’. We added covariates to the input file, specifically a ‘bimbam dosage’ file output using plink (Purcell et al. 2007; Bresadola et al. 2019). Because body mass in deer is known to be strongly influenced by age and sex (Clutton-Brock et al. 1982), we ran the BSLMM including these as additional covariates. We also included the point estimate of Q score from ADMIXTURE (see above) as an additional covariate to account for background species differences (Pallares et al. 2014). We report the results of BSLMMs run both with and without the covariates. The BSLMM was run for 25 million iterations, with a burnin of 10 million iterations, and sampled every 1000 iterations after the burnin. Convergence was confirmed using plots of the MCMC distributions of PVE, PGE, and gamma (i.e. the number of SNPs included in the sparse distribution), following (Soria-Carrasco 2019). The model was run three times to ensure that a global peak was found. To determine significance, we quantified the Posterior Inclusion Probability (PIP), and with a threshold of 0.1; those SNPs with a PIP higher than 0.1 are considered significantly associated with the phenotype (Chaves et al. 2016). We report in the main text all SNPs that we found to be significant in any of the all three runs of the model, and report the different effect sizes and PIPs from each run in Supplementary Table 1. We report exact estimates of PVE, PGE, and PIPs from the first run of the model (A in Supplementary Table 1), as all estimates were highly consistent.

To understand how genotypes for each highlighted SNP were associated with carcass mass, we used ADMIXTURE to determine the posterior population allele frequency in the parental red deer and sika populations, and to assign a ‘sika’ and a ‘red deer’ allele(s) (Alexander et al. 2009). We then plotted SNP genotypes for each sex against carcass mass after accounting for age (Wickham 2011).

### Gene Enrichment Analysis

To identify possible genes associated with carcass mass in red deer and sika, we first quantified the average linkage disequilibrium (LD) across each linkage group in each of red deer, sika and hybrids (as defined in McFarlane et al. 2020), using PLINK (Purcell et al. 2007). We used biomaRt (Durinck et al. 2005; Durinck et al. 2009) and ensembl (Yates et al. 2020) to identify genes 500kb up or downstream of the SNPs of interest, based on the high LD we expect at this range. We also used biomaRt and ensembl to infer putative function of these genes in other organisms, specifically cattle and humans. Finally, we used g:Profiler for functional gene enrichment analysis, searching for relationships between gene in predefined gene sets, where genes are categorized together based on biochemical pathways, or consistent co-expression (Subramanian et al. 2005; Raudvere et al. 2019). We compared the identified genes and associated GO terms to the databases of each cattle and humans. To account for multiple testing we used a Benjamini-Hochberg FDR, and examined each biological process (BP), molecular function (MF) and cellular component (CC) GO terms.

All data and scripts for this project can be found at https://figshare.com/projects/Admixture_mapping_reveals_loci_for_carcass_mass_in_red_deer_x_sika_hybrids_in_Kintyre_Scotland/112743.

## Results

The red deer that we sampled had an average carcass mass of 37.2kg (±17.9, females, 39.5kg± 24.2, males) while the sika weighed 19.4kg±4.3 (females, 19.3kg±9.3 males), and hybrid individuals were intermediate at 22.7kg±16.8 (females, 27.1kg±21.5 males). The substantial variation within each sex-species is due to variation in age (Figure 1).

**Figure 1:**
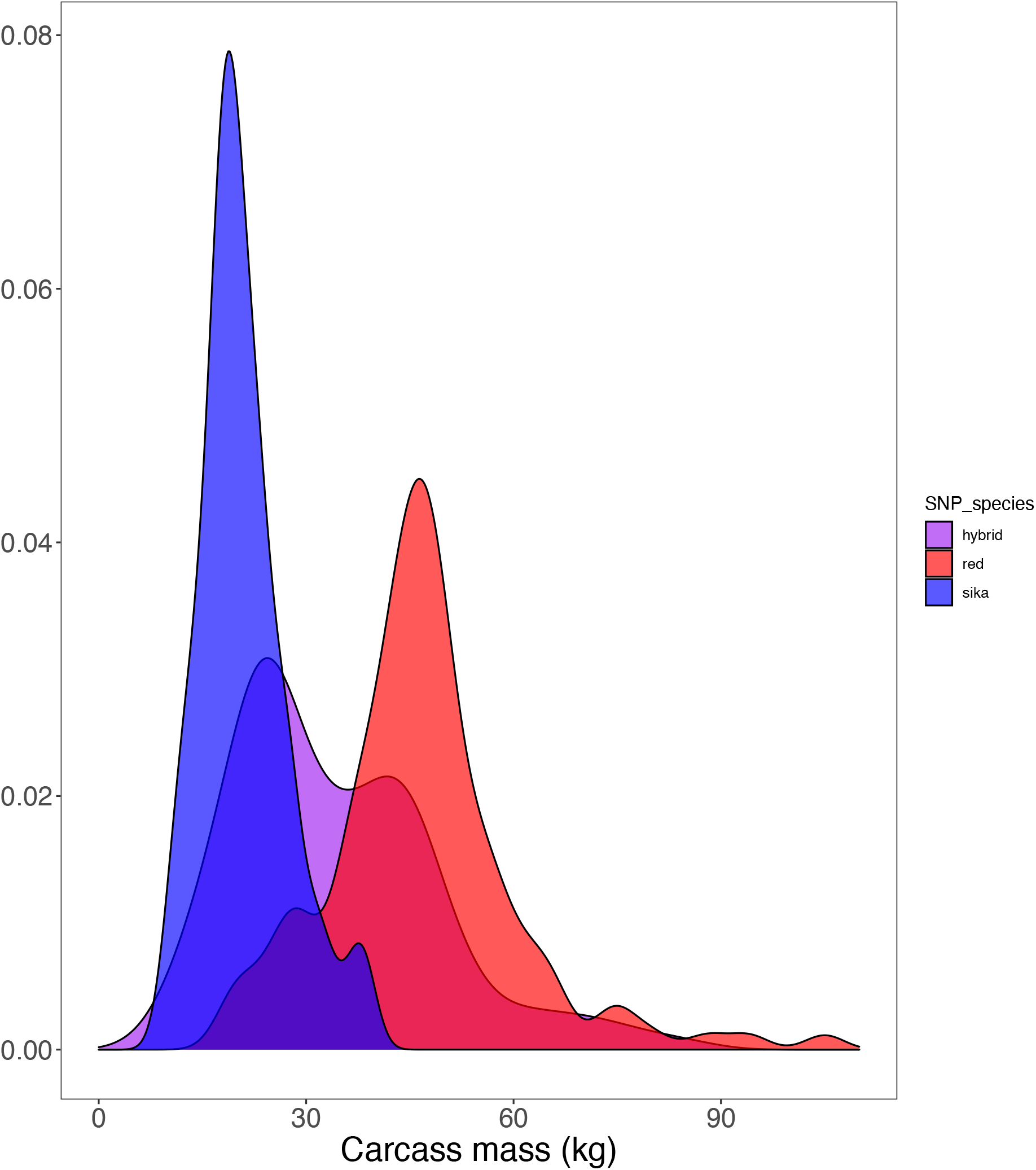
Kernal smoothed density plot of red deer (in red), sika (in blue) and hybrids (in purple) at different carcass mass in kg. Carcass mass is between 60 and 70% of live mass.

We estimated PVE, which includes sex, age, admixture proportion (Q) and the SNP effects (including the alpha matrix that included all SNP effects and the sparse matrix with the additional, large SNP effects) to explain 0.912 (0.87 – 0.95 (credible interval, CI)) of the phenotypic variance in carcass mass, while PGE, the genetic variance due to the sparse effects was 0.656 (0.22 −0.99) of. This means that 0.598 (0.19 – 0.94) of the phenotypic variance was explained by the sparse effects (i.e. those effects due to SNPs with large effects, PGE/(PVE+PGE+residual)). The sparse effects included sex and age, which both had extremely high PIP (PIPs of 1.00, 0.99 respectively) but not Q, which had a low PIP (0.0015).

The mean number of SNPs included in the sparse effects was 58.1 (6-179), but only 10 (11 in run C, Supplementary Table 1) SNPs had a PIP above the threshold of 0.1 (i.e. these SNPs were included in the sparse effect distribution at least 10% of the time). These SNPs were on linkage groups 6, 9, 11, 19, 21, 25, 28 and the X chromosome. All SNPs with a PIP higher than 0.1 were also in the 99.9^th^ percentile of effect sizes (Table 1; Figures 2 & 3), and some are clustered, for example, those on linkage group 19. One SNP on the X chromosome initially appeared to be associated with carcass mass. cela1_red_x_128791597 appeared to have a large effect size, particularly in males, but this was the result of an extremely low frequency of the minor allele associated with large carcass mass, since only one male had the relevant allele. For this reason, we believe that the effect of this SNPs is a sampling artefact, and we do not consider it any further. All other outlier SNPs had a minor allele frequency (MAF) greater than 0.15 (Table 1).

**Table 1:** SNPs that were included in the sparse distribution at least 10% of the time (i.e. PIP equal or more than 0.10). One SNP (cela1_red_3_12407635) only had a PIP above 0.1 in one of the three replicate runs of GEMMA (Supplementary Table 1). We report here the effect size (in kgs) and the posterior inclusion probability (PIP). We also note the major allele in sika and in red deer, and the allele frequency of these alleles in the parental species. While the sika alleles are nearly fixed in sika for these SNPs, red deer are polymorphic for nearly all SNPs. Finally, we report the amount (alpha estimate) and rate of (beta estimate) introgression (bgc category) for each SNP, as estimated in McFarlane et al. 2021.

| Red deer linkage group | SNP name | Effect Size | PIP | Sika Allele | Sika allele Frequency (in sika) | Red deer Allele | Red allele frequency (in red deer) | bgc category |
| --- | --- | --- | --- | --- | --- | --- | --- | --- |
| 6 | cela1_red_6_92593028 | 0.348 | 0.124 | G | 1.000 | A | 0.532 | not significant not significant |
| 9 | cela1_red_7_76865763 | 0.569 | 0.175 | A | 1.000 | G | 0.774 | not significant not significant |
| 11 | cela1_red_11_67732563 | 0.391 | 0.128 | C | 0.995 | C | 0.613 | not significant not significant |
| 19 | cela1_red_1_65547414 | 0.453 | 0.165 | A | 0.990 | G | 0.597 | not significant not significant |
| 19 | cela1_red_1_62769154 | 0.564 | 0.162 | A | 0.990 | A | 0.843 | not significant not significant |
| 20 | cela1_red_3_12407635 | 0.191 | 0.078* | G | 0.995 | G | 0.504 | not significant not significant |
| 21 | cela1_red_14_44586160 | 0.412 | 0.149 | A | 0.990 | A | 0.516 | not significant not significant |
| 25 | cela1_red_20_30400180 | 1.839 | 0.515 | C | 1.000 | A | 0.686 | not significant not significant |
| 25 | cela1_red_20_41494052 | 0.290 | 0.101 | G | 1.000 | A | 0.673 | not significant not significant |
| 28 | cela1_red_9_19132996 | 0.306 | 0.110 | A | 1.000 | A | 0.508 | not significant not significant |
| X | cela1_red_x_128791597 | 3.504 | 0.655 | G | 1.000 | A | 0.943 | not significant not significant |

**Figure 2:**
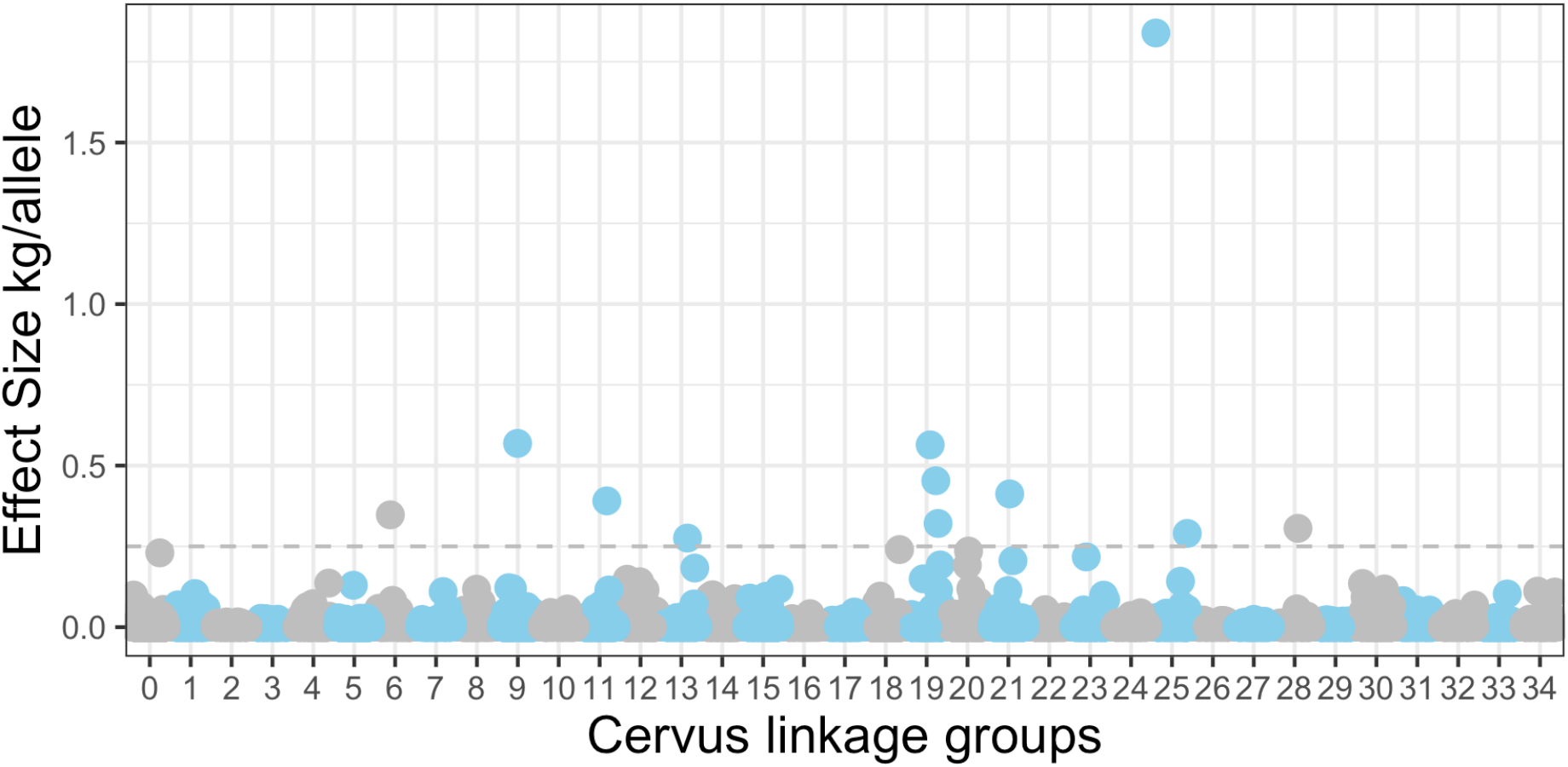
Effect size for each SNP in the sparse distribution across 34 *Cervus* linkage groups in an analysis of carcass mass, where group 0 are SNPs that are unmapped and 34 is the X chromosome. Age, sex and Q score were included in this analysis. There is one SNP on the X (chromosome 34, cela1_red_x_128791597) that is not on the plot. This SNP has an extremely high effect size, likely due to the few individuals with the minor allele (see Table 1).

**Figure 3:**
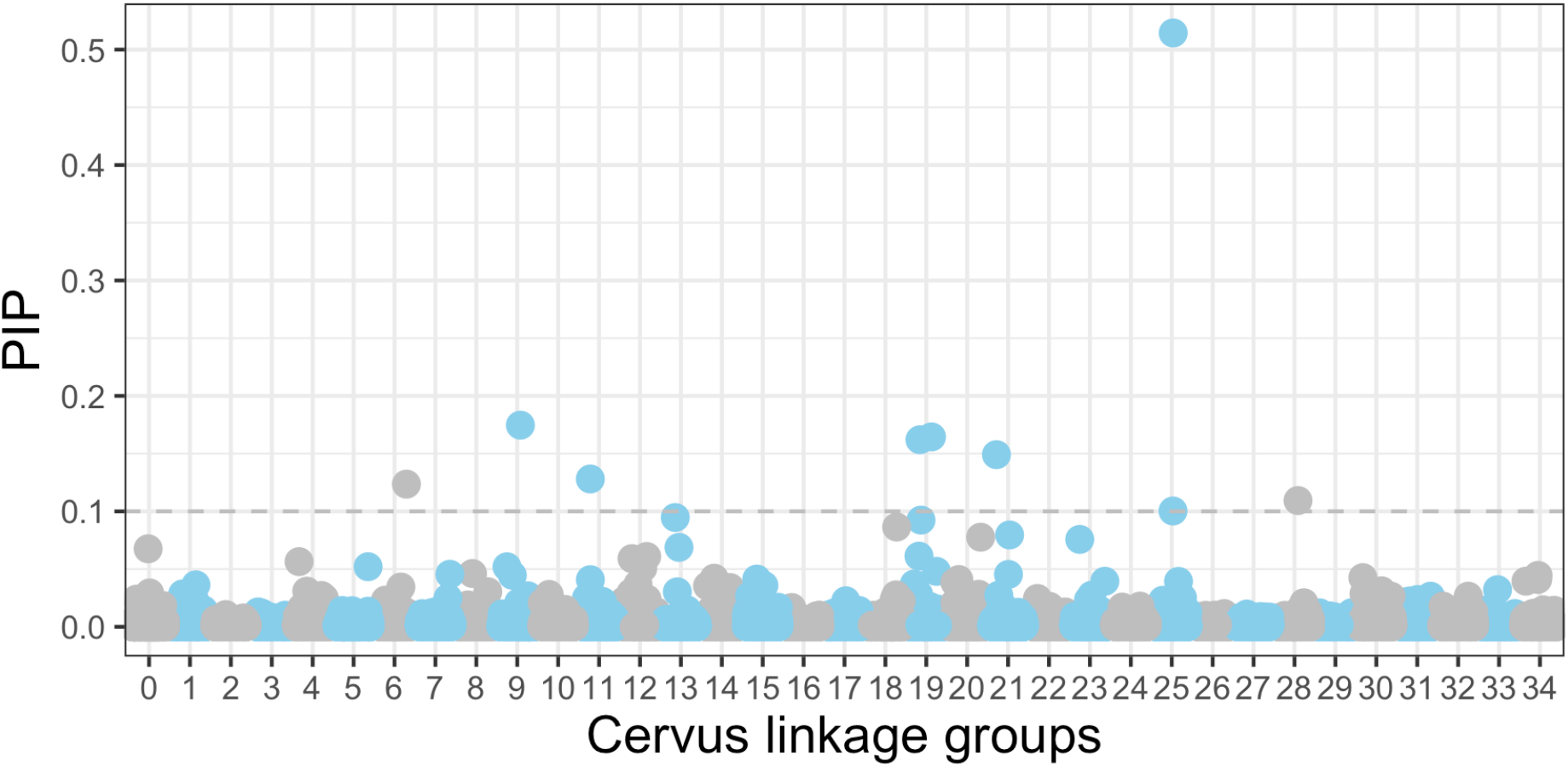
Posterior Inclusion Probability (PIP) in the sparse distribution for each SNP across 34 *Cervus* linkage groups in an analysis of carcass mass, where group 0 are SNPs that are unmapped and 34 is the X chromosome. Age, sex and Q score were included in this analysis. SNPs that were included in the sparse distribution at least 10% of the time (those above the grey dashed line) are considered significant. There is one SNP on the X (chromosome 34, cela1_red_x_128791597) that is not on the plot. This SNP has an extremely PIP, likely due to the few individuals with the minor allele (see Table 1).

To examine whether those alleles associated with small size were more prevalent in sika, we categorized alleles as sika or red deer alleles, based on posterior estimates of parental population allele frequencies from ADMIXTURE. For all SNPs significantly associated with carcass mass, the allele for small carcass mass was fixed or nearly fixed in sika, and the large carcass mass allele had a high population allele frequency in red deer (Figure 4, Table 1).

**Figure 4:**
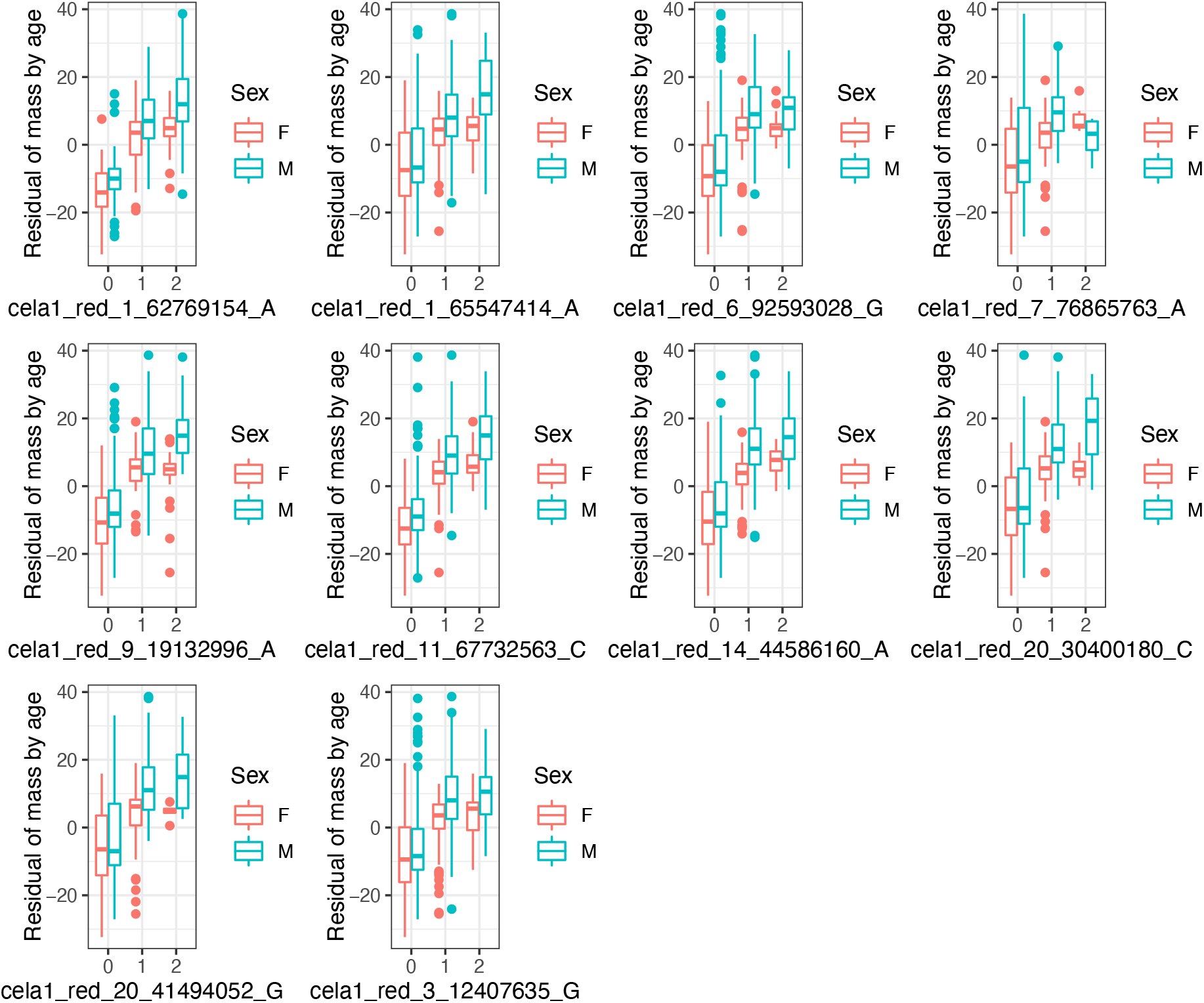
Box and whisker (central line = median, boxes 25%-75%, lines 5% - 95%) plots illustrating the relationship between the genotypes of each SNP that was included in the sparse distribution at least 10% of the time and carcass mass of each male and female deer. The x axis label notes the SNP name. ‘0’ indicates homozygotes for the sika allele, 1 heterozygotes for the sika allele and a red allele, and 2 homozygotes for the same red allele. Carcass weights have been regressed against age, and then plotted in each sex separately. A similar plot of the raw mass (instead of the residual mass) can be found in the Supplementary material (Supplementary Figure 2).

We found an average, within linkage group, LD of 0.425+/−0.26 in red deer, 0.435 +/− 0.22 in hybrid deer and 0.781+/−0.26 in sika, which varied across linkage groups (Supplementary Figure 1). From this high LD, we conservatively inferred that genes within 500Kbp of each of the significant SNPs could be related to carcass mass. We found 45 unique genes that have been named in the cattle genome (Supplementary Table 2), and 297 unique GO terms (Supplementary Table 3). We found 15 GO terms that were significantly associated via gene set analysis based on the genes that we identified (Table 3; Supplementary Table 4). We found qualitatively similar interactions when we assessed the GO terms and interactions in humans, although without any significant GO:BP interactions, and without any three-gene interactions (Supplementary Table 4).

## Discussion

Red deer and sika have a substantial phenotypic size difference, while hybrid deer are intermediate in size (Senn et al. 2010). We have identified 10 autosomal SNPs that are related to carcass mass, which are associated with seven chromosomes, 45 genes, and 297 GO terms. Our use of an anthropogenic hybrid swarm for admixture mapping has illuminated potential candidate regions in red deer and sika, which could explain variation in mass in other deer species, or even other mammalian systems.

We have identified a number of candidate regions that are associated with carcass mass in red deer, sika and their hybrids. The 10 autosomal SNPs that we have identified are all extremely invariant in sika (sika allele frequency>0.99), but polymorphic in red deer, as determined by ADMIXTURE (Alexander et al. 2009; McFarlane et al. 2020). In every case, the allele that was fixed in sika was associated with smaller size (Figure 4). Additionally, based on substantial LD across each linkage group (Table 2), we identified 40 genes that could be functionally associated with carcass mass in deer (Table S1). While none of these genes have been associated with carcass mass in white-tailed deer (Anderson et al. 2020) or cattle (cow QTL database)(Bouwman et al. 2018; Hu et al. 2019), four of them are associated with height in humans (Locke et al. 2015) and 12 are associated with body mass index (BMI) or obesity in humans (Comuzzie et al. 2012; Danjou et al. 2015; Winkler et al. 2015; Wojcik et al. 2019). Perhaps the strongest evidence we have for carcass mass QTL is where multiple adjacent SNPs indicate an effect. We found multiple SNPs on linkage groups 19 and 25, and in both cases these SNPs have large effect sizes as well as significant PIPs (Figure 2). We found 25 genes within 500000bp of the two SNPs identified on linkage group 25, and 6 of these genes were part of pre-determined gene sets, with associated GO terms (Table 2). Specifically, HBM, HBA and HBQ1 are all found on linkage group 25 and are associated with oxygen binding and transport (Table 2). Future functional work could explore if oxygen binding and transport influence growth in deer.

**Table 2:** Identified, significant gene ontology terms that are enriched across genes near the SNPs associated with carcass mass in red deer and sika when compared to the cattle genome. Possible gene ontology sources are GO:Molecular Function (GO:MF), GO:Biological Processes (GO:BP), and GO:Cellular Components (GO:CC). The association between the identified genes noted in Gene Interactions and the identified SNPs in this study can be found in Supplementary Table 2.

| source | GO Term name | GO term_id | Adjusted p_value | -Log10 Pvalue | Intersection size | Gene interactions |
| --- | --- | --- | --- | --- | --- | --- |
| GO:MF | oxygen carrier activity | GO:0005344 | 0.000 | 3.520 | 3 | HBM,HBA,HBQ1 |
| GO:MF | oxygen binding | GO:0019825 | 0.000 | 3.514 | 3 | HBM,HBA,HBQ1 |
| GO:MF | molecular carrier activity | GO:0140104 | 0.004 | 2.368 | 3 | HBM,HBA,HBQ1 |
| GO:MF | alkylbase DNA N-glycosylase activity | GO:0003905 | 0.022 | 1.652 | 1 | MPG |
| GO:MF | DNA-3-methyladenine glycosylase activity | GO:0008725 | 0.022 | 1.652 | 1 | MPG |
| GO:MF | heme binding | GO:0020037 | 0.022 | 1.652 | 3 | HBM,HBA,HBQ1 |
| GO:MF | DNA-3-methylbase glycosylase activity | GO:0043733 | 0.022 | 1.652 | 1 | MPG |
| GO:MF | DNA-7-methylguanine glycosylase activity | GO:0043916 | 0.022 | 1.652 | 1 | MPG |
| GO:MF | tetrapyrrole binding | GO:0046906 | 0.022 | 1.652 | 3 | HBM,HBA,HBQ1 |
| GO:MF | DNA-7-methyladenine glycosylase activity | GO:0052821 | 0.022 | 1.652 | 1 | MPG |
| GO:MF | DNA-3-methylguanine glycosylase activity | GO:0052822 | 0.022 | 1.652 | 1 | MPG |
| GO:MF | 2,4-dienoyl-CoA reductase (NADPH) activity | GO:0008670 | 0.041 | 1.389 | 1 | DECR2 |
| GO:BP | gas transport | GO:0015669 | 0.001 | 3.008 | 3 | HBM,HBA,HBQ1 |
| GO:BP | oxygen transport | GO:0015671 | 0.001 | 3.008 | 3 | HBM,HBA,HBQ1 |
| GO:CC | hemoglobin complex | GO:0005833 | 0.000 | 3.753 | 3 | HBM,HBA,HBQ1 |

It would be interesting to quantify selection on the specific SNPs that we have found here, to determine the potential for these genomic regions to respond to selection on body size. We have previously identified SNPs that are introgressing faster than expected from red deer into sika in our sample (McFarlane et al. 2021), but none of the SNPs associated with carcass mass are introgressing faster than the genome-wide expectation, although this doesn’t eliminate the possibility of selection for carcass mass alleles within each population. Ideally, we would measure selection on the phenotypes of hybrid individuals with a variety of genotypes to make firm statements about selection on the carcass mass loci we have identified here, and then to make predictions about the potential for adaptive introgression (Taylor and Larson 2019). However, in lieu of directly measuring fitness, admixture mapping is one piece of evidence to identify regions of the genome that are potentially contributing to introgression in hybrid systems, particularly for traits such as carcass mass which can be assumed to be under selection. It is because admixture mapping is so inherently powerful that we were able to identify SNPs explaining a substantial proportion of phenotypic and genetic variance in a quantitative trait in this wild deer system.

## Supporting information

Supplementary Table 3 Autosomal GO terms

Supplementary Table 2 Autosomal Genes

## Acknowledgements

We would like to thank Cassandre Pyne and Elizabeth Mandeville for discussions about GEMMA, Lucy Peters for help with gene identification, Forestry and Land Scotland for sample collection, especially Fraser Robertson and Kevin McKillop. We would also like to thank Helen Senn, Stephanie Smith and Rebecca Holland for the curation of samples, and for DNA extraction. Samples were genotyped at the Wellcome Trust Clinical Research Facility Genetics core, and analyses were run on Edinburgh Compute and Data Facility (ECDF; http://www.ecdf.ed.ac.uk/). This work was funded by a European Research Council Advanced Grant to JMP and a Vetenskapsrådet (Swedish Research Council) International Postdoc Fellowship to SEM.

