## Supplementary Materials for "Admixture mapping reveals loci for carcass mass in red deer x sika hybrids in Kintyre, Scotland"

**Supplementary Material:**

**Supplementary Table 1:** SNPs that were included in the sparse distribution at least 10% of the time (i.e. PIP equal or more than 0.10). One SNP (cela1_red_3_12407635) only had a PIP above 0.1 in one of the three replicate runs of GEMMA, otherwise all GEMMA estimates were highly consistent across runs A, B and C. We report here the effect size (in kgs), the posterior inclusion probability (PIP). We also note the major allele in each sika and in red deer, and the allele frequency of these alleles. While the sika alleles are nearly fixed in sika for these SNPs, red deer are polygenic for nearly all SNPs. Finally, we report the amount (alpha estimate) and rate of (beta estimate) introgression (bgc category) for each SNP, as estimated in McFarlane et al. 2021.

|  |  |  |  |  |  |  | GEMMA A | | GEMMA B | | GEMMA C | |
| --- | --- | --- | --- | --- | --- | --- | --- | --- | --- | --- | --- | --- |
| Red deer linkage group | SNP name | Sika Allele | Sika allele Frequency (in Sika) | Red deer Allele | Red Allele Frequency (in Red deer) | bgc category | Effect Size | PIP | Effect Size | PIP | Effect Size | PIP |
| 34 | cela1_red_x_128791597 | G | 1.000 | A | 0.943 | not significant not significant | 3.504 | 0.655 | 3.635 | 0.665 | 3.782 | 0.679 |
| 25 | cela1_red_20_30400180 | C | 1.000 | A | 0.686 | not significant not significant | 1.839 | 0.515 | 1.814 | 0.505 | 1.784 | 0.487 |
| 9 | cela1_red_7_76865763 | A | 1.000 | G | 0.774 | not significant not significant | 0.569 | 0.175 | 0.687 | 0.213 | 0.725 | 0.217 |
| 19 | cela1_red_1_65547414 | A | 0.990 | G | 0.597 | not significant not significant | 0.453 | 0.165 | 0.423 | 0.152 | 0.396 | 0.143 |
| 19 | cela1_red_1_62769154 | A | 0.990 | A | 0.843 | not significant not significant | 0.564 | 0.162 | 0.569 | 0.154 | 0.409 | 0.120 |
| 21 | cela1_red_14_44586160 | A | 0.990 | A | 0.516 | not significant not significant | 0.412 | 0.149 | 0.406 | 0.147 | 0.305 | 0.112 |
| 11 | cela1_red_11_67732563 | C | 0.995 | C | 0.613 | not significant not significant | 0.391 | 0.128 | 0.343 | 0.115 | 0.323 | 0.109 |
| 6 | cela1_red_6_92593028 | G | 1.000 | A | 0.532 | not significant not significant | 0.348 | 0.124 | 0.314 | 0.106 | 0.279 | 0.098 |
| 28 | cela1_red_9_19132996 | A | 1.000 | A | 0.508 | not significant not significant | 0.306 | 0.110 | 0.280 | 0.102 | 0.359 | 0.132 |
| 25 | cela1_red_20_41494052 | G | 1.000 | A | 0.673 | not significant not significant | 0.290 | 0.101 | 0.294 | 0.101 | 0.263 | 0.087 |
| 20 | cela1_red_3_12407635* | G | 0.995 | G | 0.504 | not significant not significant | 0.191 | 0.078 | 0.195 | 0.080 | 0.246 | 0.101 |

**Supplementary Table 2:** Autosomal SNPs associated with carcass weight in red deer, sika and hybrid deer in Kintyre, Scotland, and named genes within 500,000 BP of each SNP. The second row denotes the red deer linkage group each SNP is found on (Johnston et al. 2017).

This table is an excel file named: Supplementary Table 2 – Autosomal Genes

**Supplementary Table 3:** Autosomal SNPs associated with carcass weight in red deer, sika and hybrid deer in Kintyre, Scotland, and Gene Ontogeny (GO) terms associated with the genes near each SNP (see Supplementary Table 2). The second row denotes the red deer linkage group each SNP is found on.

This table is an excel file named: Supplementary Table 3 – Autosomal GO-terms

**Supplementary Table 4**: Identified, significant gene ontology terms that are enriched across genes near the SNPs associated with carcass mass in red deer and sika when compared to the human genome. Possible gene ontology sources are GO:Molecular Function (GO:MF), and GO:Cellular Components (GO:CC). No interactions were found in the GO:Biological Processes source (GO:BP). The association between the identified genes noted in Gene Interactions and the identified SNPs in this study can be found in Supplementary Table 3.

| **source** | **GO Term name** | **GO term_id** | **Adjusted p_value** | **-Log10 Pvalue** | **Intersection size** | **Gene interactions** |
| --- | --- | --- | --- | --- | --- | --- |
| GO:MF | haptoglobin binding | GO:0031720 | 0.029 | 1.532 | 2 | HBM,HBQ1 |
| GO:MF | oxygen carrier activity | GO:0005344 | 0.034 | 1.468 | 2 | HBM,HBQ1 |
| GO:MF | alkylbase DNA N-glycosylase activity | GO:0003905 | 0.040 | 1.396 | 1 | MPG |
| GO:MF | interleukin-9 receptor activity | GO:0004919 | 0.040 | 1.396 | 1 | IL9R |
| GO:MF | DNA-3-methyladenine glycosylase activity | GO:0008725 | 0.040 | 1.396 | 1 | MPG |
| GO:MF | DNA-3-methylbase glycosylase activity | GO:0043733 | 0.040 | 1.396 | 1 | MPG |
| GO:MF | DNA-7-methylguanine glycosylase activity | GO:0043916 | 0.040 | 1.396 | 1 | MPG |
| GO:MF | DNA-7-methyladenine glycosylase activity | GO:0052821 | 0.040 | 1.396 | 1 | MPG |
| GO:MF | DNA-3-methylguanine glycosylase activity | GO:0052822 | 0.040 | 1.396 | 1 | MPG |
| GO:MF | oxygen binding | GO:0019825 | 0.047 | 1.331 | 2 | HBM,HBQ1 |
| GO:CC | hemoglobin complex | GO:0005833 | 0.021 | 1.677 | 2 | HBM,HBQ1 |
| GO:CC | haptoglobin-hemoglobin complex | GO:0031838 | 0.021 | 1.677 | 2 | HBM,HBQ1 |
| GO:CC | U2-type prespliceosome | GO:0071004 | 0.021 | 1.677 | 2 | SNRPN,LUC7L |
| GO:CC | prespliceosome | GO:0071010 | 0.021 | 1.677 | 2 | SNRPN,LUC7L |
| GO:CC | U1 snRNP | GO:0005685 | 0.024 | 1.622 | 2 | LUC7L,SNRPN |

**Supplementary Figure 1 :** Mean linkage disequilibrium for SNPs within 10kb for each red deer, hybrid deer and sika from Kintyre. The sika LD is very high, likely because sika alleles are near fixation at most loci. Specifically, 33 755 SNPs have a minor allele frequency lower than 0.05 in pure sika individuals.

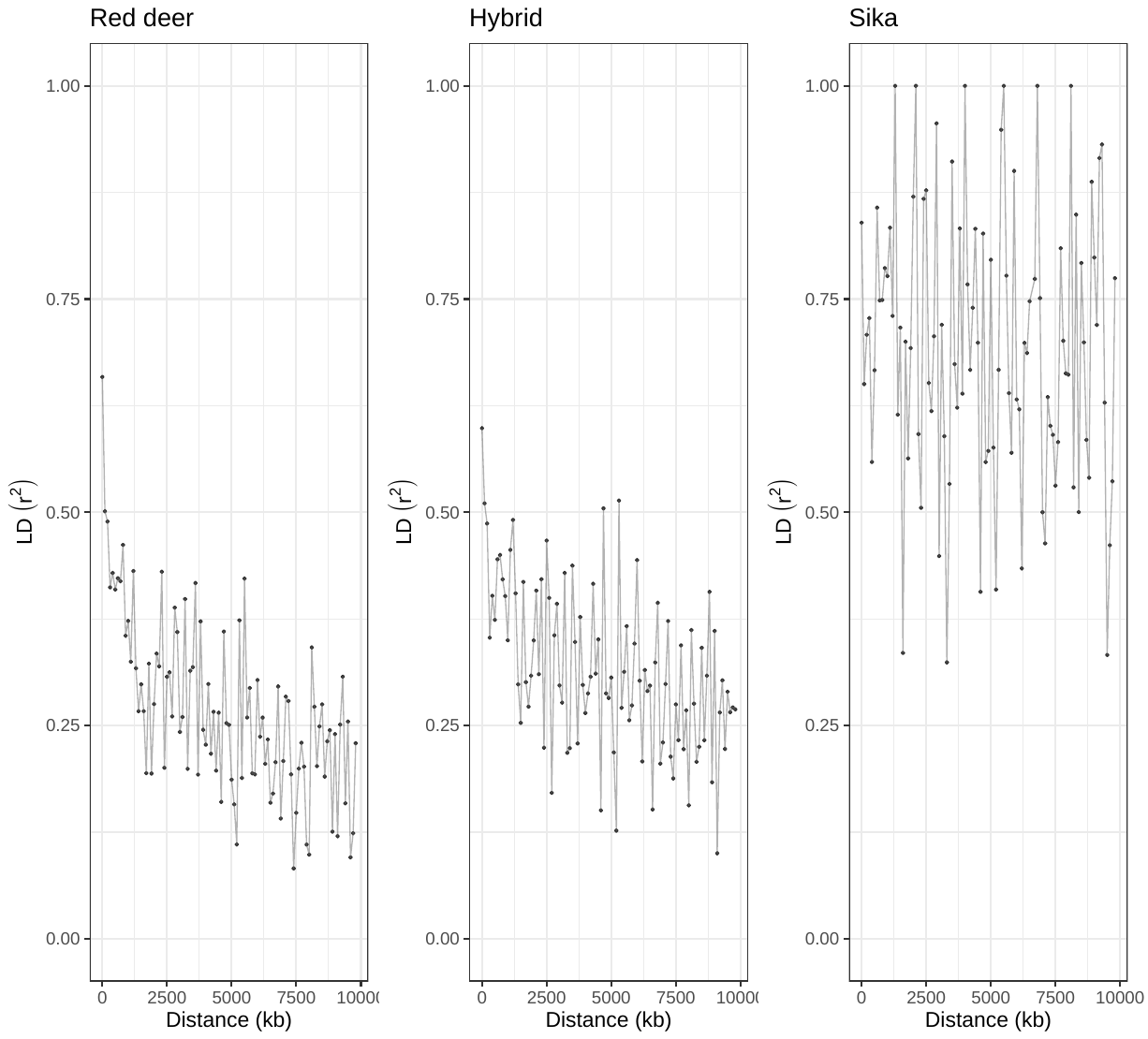

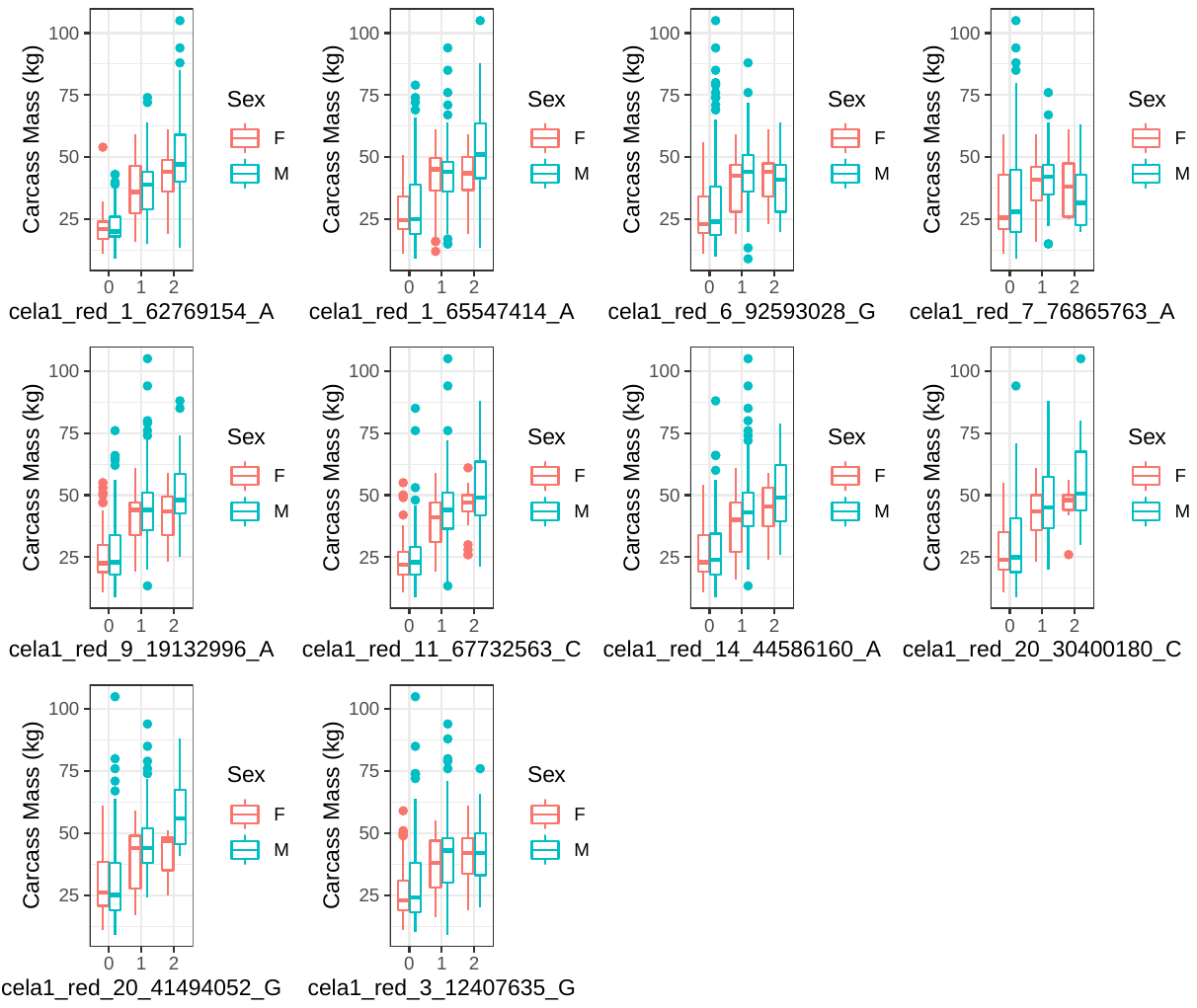

**Supplementary Figure 2:** Box and whisker (central line = median, boxes 25%-75%, lines 5% - 95%) plots illustrating the relationship between the genotypes of each SNP that was included in the sparse distribution at least 10% of the time and carcass mass of each male and female deer. The x axis label notes both the SNP name, and the allele associated with the minor allele (i.e. 0), which is the ‘sika’ allele in all cases. This plot is on the raw data, and does not account for age, an additional known covariate of carcass mass.
